# Subcortical volumes, frontal cortical thickness, and pro-inflammatory cytokines in schizophrenia versus methamphetamine-induced psychosis

**DOI:** 10.1101/2023.09.18.558193

**Authors:** Lauren Blake, Kimberley Clare Williams, Anne Alex Uhlmann, Henk Temmingh, Antoinette Burger, Dan Joseph Stein, Petrus J. W. Naudé

## Abstract

**Background and Hypothesis:** Clinical features of schizophrenia and methamphetamine-induced psychosis (MAP) have significant overlap. Schizophrenia is associated with alterations in subcortical volumes and cortical thickness and pro-inflammatory cytokines, that may correlate with clinical features. However, analogous work on MAP is lacking. We hypothesise decreased frontal cortical thickness, amygdala and hippocampal volumes in schizophrenia and MAP compared to controls, with increased basal ganglia in schizophrenia compared to MAP.

**Study Design:** Participants included n=36 schizophrenia, n=27 MAP and 32 age- and sex-matched healthy controls. Diagnosis and symptom severity were determined with the Structured Clinical Interview for Axis I Disorders, and the Positive and Negative Syndrome Scale, respectively. Structural T1-weighted images were acquired using a 3-Tesla magnetic resonance imaging scanner. Serum peripheral cytokine concentrations were measured using a multiplex bead array.

**Study Results:** Schizophrenia and MAP participants showed decreased left amygdala volumes and frontal cortical thickness compared to controls. Schizophrenia participants had increased bilateral caudate, putamen, and nucleus accumbens (NAcc) volumes compared to controls, and greater right globus pallidus and NAcc volumes compared to MAP. Decreased left amygdala volumes and frontal cortical thickness were significantly associated with longer illness duration in schizophrenia and methamphetamine use in MAP. No significant differences were found in cytokine levels between groups or associations with neuroimaging measures.

**Conclusions:** This study highlights overlaps and differences in the psychobiology of schizophrenia and MAP.The novel discovery of increased globus pallidus and NAcc volumes in schizophrenia compared with MAP may show important distinctions in the neurobiology between these two conditions.

## 1. Introduction

There is considerable overlap in the clinical features of schizophrenia and methamphetamine-induced psychosis (MAP). Both schizophrenia and MAP are characterized by positive symptoms of psychosis. Thus, it has been suggested that MAP may serve as an important model for understanding the neurobiology of schizophrenia. Nevertheless, few studies have directly compared the biosignatures of schizophrenia and MAP.

Structural magnetic resonance imaging (MRI) studies have shown differences in subcortical gray matter volumes, in schizophrenia, which have been associated with clinical symptom profiles ^1–5^. In schizophrenia, decreased hippocampal volume has been associated with negative symptoms and hallucinations ^4,5^, with decreased amygdala-hippocampal complex associated with negative symptoms of psychosis ^5^. Furthermore, increased basal ganglia volumes have been consistently observed in schizophrenia compared to healthy controls ^6–8^. Alterations in basal ganglia volumes have also been associated with clinical symptom profiles ^1–5^ including positive and negative symptoms ^9^. Differences in cortical thickness of the frontal and temporal lobes have also been observed in individuals with schizophrenia ^10–15^, with the frontal cortex being most affected ^15^. While structural neuroimaging studies of MAP are limited, there is some evidence of increased cortical thickness in MAP compared to individuals with methamphetamine dependence ^16^ and decreased hippocampal and amygdala volumes in MAP compared to healthy controls ^17^. However, no studies have directly compared subcortical volumes and cortical thickness in schizophrenia to that in MAP.

Schizophrenia is further characterized by a number of alterations in immune biomarkers (Prestwood et al., 2021). Alterations of peripheral blood inflammatory cytokines (e.g. TNF)-α, IL-1β, IL-6, IL-8, IL-10, IL-12, and IFN-γ) have been associated with clinical outcomes in schizophrenia ^18,19^. However, few studies have investigated immune markers in MAP, and none have directly compared immune markers in schizophrenia to those in MAP. Given findings of associations between neuroimaging measures and cytokine levels in schizophrenia ^20^, and evidence of neuro-inflammation in laboratory models of MA use ^21,22^, investigation on associations between neuroimaging measures and cytokines in MAP can advance our understanding on the potential functions of neuro-inflammation in schizophrenia and MAP.

This study aims to compare subcortical volumes, cortical thickness, and cytokine levels in individuals with schizophrenia, MAP, and healthy controls. We also assessed associations of neuroimaging measures with illness duration, symptom severity, and pro-inflammatory cytokines in schizophrenia and MAP, and with duration of methamphetamine use in MAP.

## 2. Materials and methods

### 2.1. Participants

In this exploratory study, participants between the ages of 20-40 years, were recruited from Valkenberg Hospital and Groote Schuur Hospital in Cape Town, and included 36 (8 females: 28 males) participants with schizophrenia, 27 (6 females: 21 males) participants with MAP and 32 (13 females: 19 males) healthy controls, matched for age and sex, were recruited from the general population in Cape Town.

Individuals with schizophrenia and MAP had clinical diagnosis confirmed with the Structured Clinical Interview for DSM-IV (SCID-I), and symptom severity measured with the Positive and Negative Syndrome Scale (PANSS) ^23^. The SCID-I was also given to healthy controls to include only those without Axis-I disorders. Participants were excluded if they had a history of 1) chronic medical illness that required medical care or prescription medication, including severe brain trauma and intellectual disability; 2) were recently or currently pregnant or lactating; 3) were unable to undergo MRI because of metal implants, claustrophobia, or other reasons; 4) substance dependence other than nicotine dependence.

Written consent was obtained from all participants included in the study. The work adhered to human research guidelines as stipulated in the Declaration of Helsinki ^24^. The protocol underwent ethical review and approval with the University of Cape Town’s Human Research Ethics Committee (UCT HSF HREC: 413/2016).

### 2.2. Imaging procedures

Structural magnetic resonance imaging (sMRI) was conducted in the morning on the second day of the study at the Cape Universities Body Imaging Centre (CUBIC), using a 3T Siemens Skyra MRI scanner (Siemens Medical Systems, Erlangen, Germany). A 32-channel head coil was used. A high resolution, T1-weighted scans were acquired with a multi-echo MPRAGE structural scan with the following scan parameters: TR= 2530ms; graded TE= 1.69, 3.55, 5.41 and 7.27ms; flip angle= 7°; field of view (FOV)= 256mm; voxel size = 1x1x1mm; 6:39min.

### 2.3. Image processing

Processing of all MRIs was performed using FreeSurfer v6.0 ^25,26^ (http://surfer.nmr.mgh.harvard.edu/) using the Desikan-Killiany atlas, for cortical reconstruction and volume segmentation ^27^. The processing pipeline included motion correction, skull-stripping, talairach transformation, segmentation of subcortical white matter and deep grey matter volumetric structures, intensity normalization, tessellation of the grey matter/white matter boundary, automated topology correction, and surface deformation and registration to the Desikan-Killiany atlas ^27^. Reconstruction of the grey and white matter boundary and pial surface was performed to extract cortical thickness measurements ^28–30^. Data and statistical summaries were generated using ENIGMA protocols (http://enigma.ini.usc.edu/protocols/imaging-protocols) and RStudio v 1.2.5054 ^31^.

### 2.4. Inflammatory markers

Blood samples were collected via venepuncture into serum tubes in the morning of the first day of the study. The tubes were kept at room temperature at for approximately 30 minutes to allow clotting and were subsequently centrifuged at 1,800 × g for 10 min. The separated serum was then aliquoted into cryovials and stored at -80 °C until analyses. The Luminex Milliplex^®^ MAP human cytokine / chemokine magnetic bead panel (HSTCMAG28SK07; Merck) was used to detect cytokines; TNF-*α*, IL-1*β*, IL-6, IL- 8, IL-10, IL-12p, and IFN-*γ*. The intra-assay coefficients of variation for each cytokine were all below 14%. All plates were analysed on the same day. Serum measures for IL-6 were below the detectable limit and were consequently excluded from statistical analysis.

### 2.5. Statistical analyses

*A priori* power analysis, to determine the minimum required sample size (n) required, was conducted with G*Power 3 ^32^ as a function of the required power level (1 – β error probability = 0.8), the pre-specified significance level (α = 0.05), and the population effect size (d = 0.5) with an allocation ratio of N2/N1 = 1. The *a priori* power analysis yielded a required total sample size of n = 102, with statistical power of 0.81.

To compare brain measures across schizophrenia (n = 36), MAP (n = 27), and healthy controls (n = 32), we employed a multiple analysis of covariance (MANCOVA) for all brain regions, with intracranial volume (ICV) ^33–35^, sex, and age ^36–38^ as covariates for subcortical volume measurements and, sex and age as covariates for frontal cortical thickness measurements. Thereafter, a multiple analysis of variance (MANOVA) for parametric data and Kruskal Wallis analysis of variance (ANOVA) for non-parametric data to determine whether there were group differences. MANOVA was followed by Bonferroni post-hoc tests to determine which groups differed from each other. Effect sizes for group comparisons were calculated using partial eta squared for all parametric ANOVAs and eta squared for Kruskal Wallis ANOVAs at a confidence interval of 90% ^39^. Spearman’s Rank correlations (for non-normally distributed data) and Pearson’s correlations (normally distributed data) were performed within each group to determine associations of neuroimaging measures with symptom severity, illness duration, duration of methamphetamine use and cytokine concentrations. Significance levels were set at Rho of ± 0.5 and *p* < 0.01 (where the critical value for r = 0.449 when *p* = 0.01) to reduce Type I Error and increase stringency and strength of the associations between variables.

In a sensitivity analysis to determine whether there were differences between the schizophrenia participants who reported methamphetamine (n = 14) use to those that did not report methamphetamine use (n = 22), we employed t-tests for parametric brain regions and Mann Whitney U tests for non-parametric brain regions (Supplementary Tables 1, 2 and 3). Effect sizes were calculated using Cohen’s D for parametric data at a 95% confidence interval and biserial correlations *r* at *p* < 0.05.

## 3. Results

### 3.1. Demographic and clinical variables

Table 1 shows that there were no significant group differences for age and sex, lower levels of school education were found in schizophrenia (H_(2,94)_= 19.827, *p* = 0.002) and in MAP (H_(2,94)_= 19.827, *p* = 0.00046), compared to healthy controls. Duration of illness was significantly longer in schizophrenia compared to MAP (*z* = 3.271, *p* = 0.0008) and the number of psychotic episodes were greater in schizophrenia than MAP (*p* = 0.038). Significant differences were found for the PANSS positive (H_(2,92)_= 40.21, *p* < 0.0001), negative (H_(2,92)_= 43.86, *p* < 0.0001), general (H_(2,92)_= 32.99, *p* < 0.0001) and total scores (H_(2,92)_= 46.64, *p* < 0.0001), with significant differences found between controls and schizophrenia (*p* < 0.0001) and, controls and MAP (*p* ≤ 0.00012).

**Table 1:**
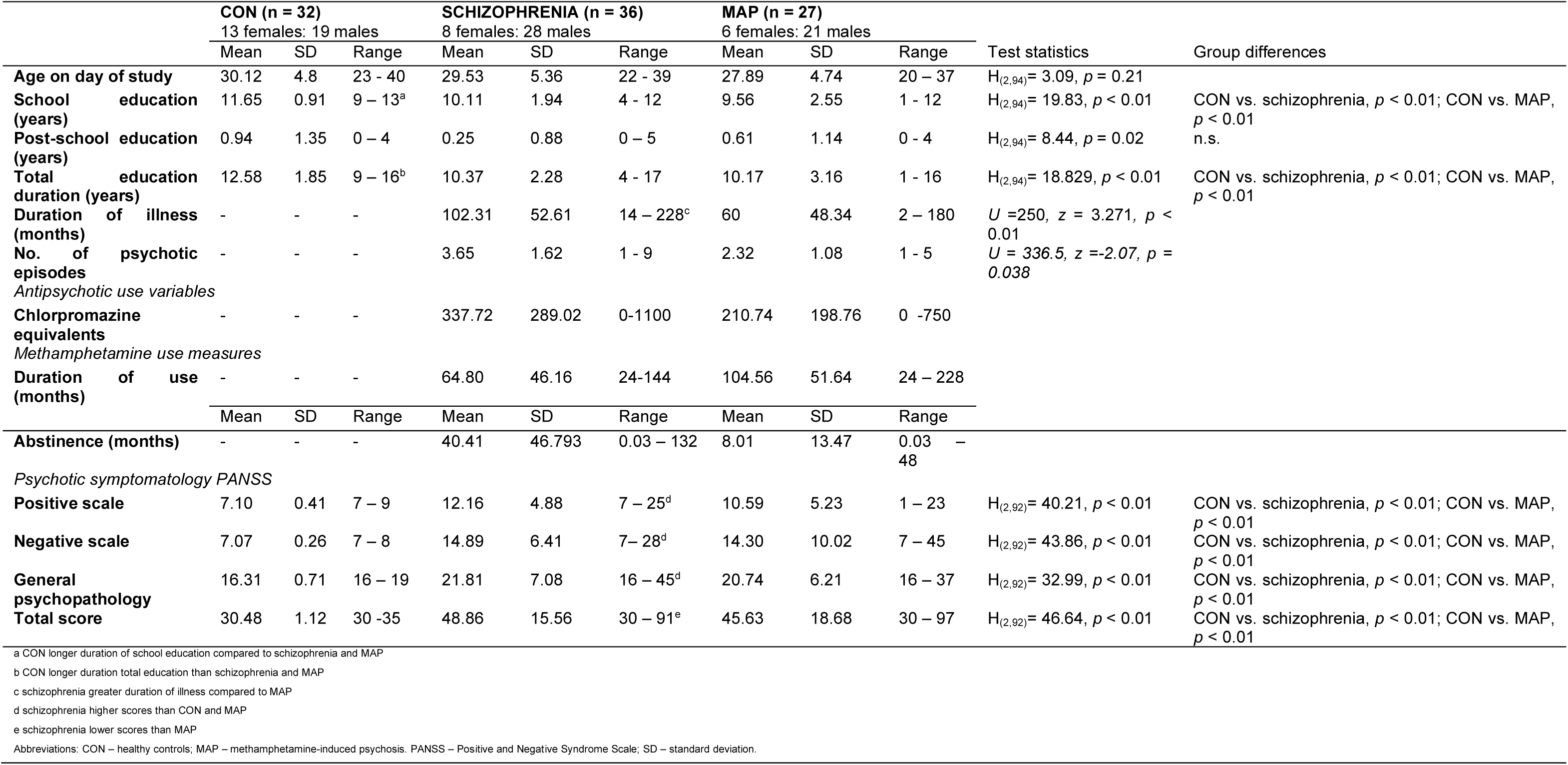
Demographics, duration of illness, clinical scales and methamphetamine use variables for control, schizophrenia and methamphetamine-induced psychotic groups.

### 3.2. Subcortical volumes

There were decreased left amygdala volumes in schizophrenia (F_(2,91)_ = 6.37, *p* = 0.012) and in MAP (F_(2,91)_ = 6.37, *p* = 0.0054) compared to healthy controls (Table 2). There were increased bilateral caudate (F_(2,91)_ = 4.11, *p* = 0.03; F_(2,91)_ = 3.95, *p* = 0.05), putamen (F_(2,91)_ = 6.85, *p* = 0.016; F_(2,91)_ = 5.27, *p* = 0.0072) and NAcc (F_(2,91)_ = 18.69, *p* < 0.0001; F_(2,91)_ = 7.83, *p* = 0.014) volumes in schizophrenia compared to healthy controls. Decreased bilateral NAcc (F_(2,91)_ = 18.69; *p* < 0.0001; F_(2,91)_ = 7.83, *p* = 0.0017) and right globus pallidus (F_(2,91)_ = 3.81, *p* = 0.021) volumes were observed in MAP compared to schizophrenia (Table 2).

**Table 2:** Subcortical volume group differences across schizophrenia, methamphetamine-induced psychosis and health controls.

| Regions | All groups |  |  | SCZ-CON | MAP-CON | MAP-SCZ |
| --- | --- | --- | --- | --- | --- | --- |
| | Effect size<br>$\eta p^2$ | CI: 90% | *p* | *p* | *p* | *p* |
| Left hippocampus | 0.012 | [0, 0.056] | 0.57 | n.s. | n.s. | n.s. |
| Right hippocampus | 0.051 | [0, 0.13] | 0.090 | n.s. | n.s. | n.s. |
| Left amygdala | 0.012 | [0.028, 0.22] | 0.0026 | 0.012 | 0.054 | n.s. |
| Right amygdala | 0.029 | [0, 0.091] | 0.26 | n.s. | n.s. | n.s. |
| Left caudate | 0.082 | [0.0075, 0.17] | 0.020 | 0.03 | n.s. | n.s. |
| Right caudate | 0.08 | [0.0062, 0.17] | 0.023 | 0.05 | n.s. | n.s. |
| Left putamen | 0.13 | [0.033, 0.23] | 0.0017 | 0.016 | n.s. | n.s. |
| Right putamen | 0.10 | [0.017, 0.2] | 0.0068 | 0.0072 | n.s. | n.s. |
| Left globus pallidus | 0.035 | [0, 0.1] | 0.19 | n.s. | n.s. | n.s. |
| Right globus pallidus | 0.076 | [0.052, 0.16] | 0.026 | n.s. | n.s. | 0.021 |
| Left nucleus accumbens | 0.29 | [0.16, 0.39] | < 0.0001 | < 0.0001 | n.s. | < 0.0001 |
| Right nucleus accumbens | 0.14 | [0.043, 0.25] | 0.0007 | 0.014 | n.s. | 0.0017 |

Effect size: for all group differences. \*MANOVA ( $F(2, 91)$ ) was performed for subcortical volumes in methamphetamine-induced psychosis (MAP), schizophrenia (SCZ) and healthy controls (CON) groups, with *p*-values of <0.05 considered statistically significant. n.s : non-significant. Effect sizes: partial eta squared (Confidence Interval: 90%)

No significant differences in subcortical volumes were observed between the schizophrenia group which reported methamphetamine use and the schizophrenia group who did not report methamphetamine use (Supplementary Table 1).

### 3.3 Frontal cortical thickness

There was decreased frontal cortical thickness in schizophrenia compared to healthy controls for all frontal regions (F_(2,91)_ > 10.5, *p* < 0.0004) except for the right lateral orbitofrontal, left and right medial orbitofrontal and right rostral middle frontal regions (Table 3).

**Table 3:**
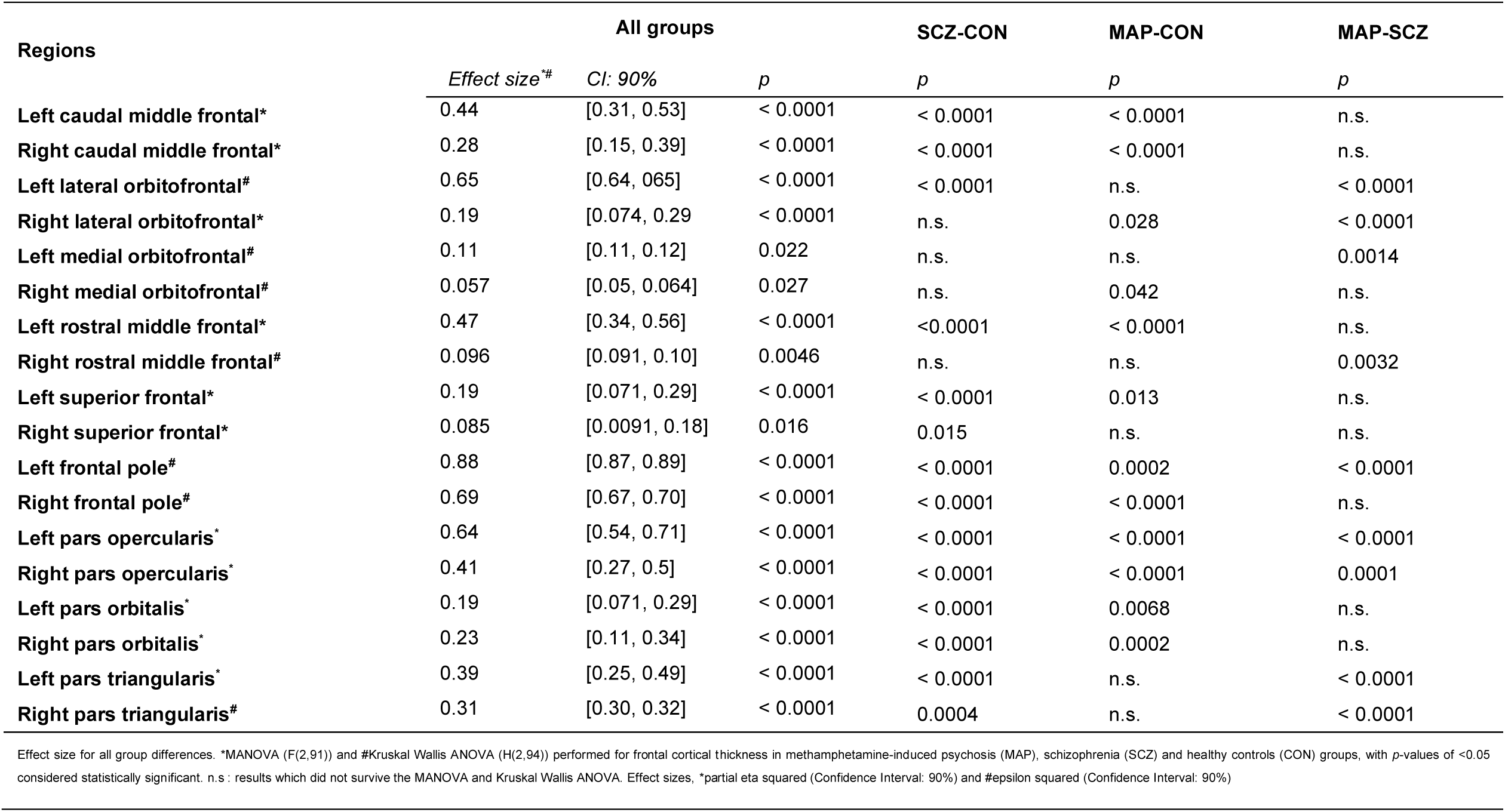
Frontal cortical thickness group differences.

| Regions | All groups |  |  | SCZ-CON | MAP-CON | MAP-SCZ |
| --- | --- | --- | --- | --- | --- | --- |
|  | Effect size*<sup>#</sup> | CI: 90% | p | p | p | p |
| Left caudal middle frontal* | 0.44 | [0.31, 0.53] | < 0.0001 | < 0.0001 | < 0.0001 | n.s. |
| Right caudal middle frontal* | 0.28 | [0.15, 0.39] | < 0.0001 | < 0.0001 | < 0.0001 | n.s. |
| Left lateral orbitofrontal<sup>#</sup> | 0.65 | [0.64, 0.65] | < 0.0001 | < 0.0001 | n.s. | < 0.0001 |
| Right lateral orbitofrontal* | 0.19 | [0.074, 0.29] | < 0.0001 | n.s. | 0.028 | < 0.0001 |
| Left medial orbitofrontal<sup>#</sup> | 0.11 | [0.11, 0.12] | 0.022 | n.s. | n.s. | 0.0014 |
| Right medial orbitofrontal<sup>#</sup> | 0.057 | [0.05, 0.064] | 0.027 | n.s. | 0.042 | n.s. |
| Left rostral middle frontal* | 0.47 | [0.34, 0.56] | < 0.0001 | < 0.0001 | < 0.0001 | n.s. |
| Right rostral middle frontal<sup>#</sup> | 0.096 | [0.091, 0.10] | 0.0046 | n.s. | n.s. | 0.0032 |
| Left superior frontal* | 0.19 | [0.071, 0.29] | < 0.0001 | < 0.0001 | 0.013 | n.s. |
| Right superior frontal* | 0.085 | [0.0091, 0.18] | 0.016 | 0.015 | n.s. | n.s. |
| Left frontal pole<sup>#</sup> | 0.88 | [0.87, 0.89] | < 0.0001 | < 0.0001 | 0.0002 | < 0.0001 |
| Right frontal pole<sup>#</sup> | 0.69 | [0.67, 0.70] | < 0.0001 | < 0.0001 | < 0.0001 | n.s. |
| Left pars opercularis* | 0.64 | [0.54, 0.71] | < 0.0001 | < 0.0001 | < 0.0001 | < 0.0001 |
| Right pars opercularis* | 0.41 | [0.27, 0.5] | < 0.0001 | < 0.0001 | < 0.0001 | 0.0001 |
| Left pars orbitalis* | 0.19 | [0.071, 0.29] | < 0.0001 | < 0.0001 | 0.0068 | n.s. |
| Right pars orbitalis* | 0.23 | [0.11, 0.34] | < 0.0001 | < 0.0001 | 0.0002 | n.s. |
| Left pars triangularis* | 0.39 | [0.25, 0.49] | < 0.0001 | < 0.0001 | n.s. | < 0.0001 |
| Right pars triangularis<sup>#</sup> | 0.31 | [0.30, 0.32] | < 0.0001 | 0.0004 | n.s. | < 0.0001 |

Effect size for all group differences. \*MANOVA (F(2,91)) and #Kruskal Wallis ANOVA (H(2,94)) performed for frontal cortical thickness in methamphetamine-induced psychosis (MAP), schizophrenia (SCZ) and healthy controls (CON) groups, with p-values of <0.05 considered statistically significant. n.s.: results which did not survive the MANOVA and Kruskal Wallis ANOVA. Effect sizes, \*partial eta squared (Confidence Interval: 90%) and #epsilon squared (Confidence Interval: 90%)

Decreased frontal cortical thickness in MAP compared to healthy controls for all frontal regions (All, F_(2,91)_ > 10.5, *p* < 0.013) was found except for the left lateral orbitofrontal, medial orbitofrontal, right rostral middle frontal and superior frontal and the left and right pars triangularis, and increased cortical thickness in MAP compared to healthy controls in the right medial orbitofrontal (H_(2,94)_ = 7.21, *p* = 0.042) and right lateral orbitofrontal (F_(2,91)_ = 10.7, *p* = 0.028).

Increased frontal cortical thickness was observed in MAP compared to schizophrenia for the right lateral orbitofrontal (F_(2,91)_ = 10.7, *p* < 0.0001 ), left pars opercularis (F_(2,91) )_ = 82.5, *p* < 0.0001), left pars triangularis (F_(2,91)_ = 29.2, *p* < 0.0001), left lateral orbitofrontal (H_(2,94)_ = 60.76, *p* < 0.0001), left medial orbitofrontal (H_(2,94)_ = 12.22, *p* = 0.0014), left frontal pole (H_(2,94)_ = 82.30, *p* < 0.0001 ) and left and right pars triangularis (F_(2,91)_ = 29.2, *p* < 0.0001). There was increased frontal cortical thickness was observed in schizophrenia compared to MAP for the right rostral middle frontal (H_(2,94)_ = 10.78, *p* = 0.0032) and the right pars opercularis (F_(2,91)_ = 31.26, *p* = 0.0001).

Increased frontal cortical thickness was observed in schizophrenia group which used methamphetamine for the left lateral orbitofrontal (*U* = 64.5, *p* = 0.039) and right medial orbitofrontal (*U* = 67.5, *p* = 0.05) compared to the schizophrenia group who reported no methamphetamine use. Decreased frontal cortical thickness was observed in the schizophrenia group who reported methamphetamine use for the left frontal pole (*U* = 178.5, *p* = 0.039) compared to the schizophrenia group who reported no methamphetamine use (Supplementary Tables 2 and 3).

### 3.4. Cytokines

There were no significant group differences in cytokine concentrations for IFN-γ (H_2,94_ = 2.81; *p* = 0.245), IL-10 (H_2,94_ = 0.93, *p* = 0.627), IL-12 (H_2,94_ = 4.90, *p* = 0.0862), IL-1β (H_2,94_ = 1.27, *p* = 0.531), IL-8 (H_2,94_ = 0.23; *p* =0.892) and TNF-α (H_2,94_ = 1.01, *p* = 0.605).

### 3.5. Associations of neuroimaging measures with pro-inflammatory cytokines, illness duration and symptom severity

There were no significant associations between neuroimaging measures and either symptom severity with cytokine concentrations.

In schizophrenia, longer illness duration was associated with decreased left amygdala volumes (R_(n=36)_ = -0.43, *p* = 0.0088), and decreased frontal cortical thickness (All, R_(n=36)_ < -0.51, *p* < 0.0013), except for the left frontal pole (R_(n=36)_ = 0.48, *p* = 0.0029) and right frontal pole (R_(n=36)_= 0.37, *p* = 0.025).

In MAP, increased duration of methamphetamine use was associated with decreased left amygdala (r_(n=27)_ = -0.51, *p* = 0.0069) volumes, left lateral orbitofrontal (R_(n=27)_ = -0.54, *p* = 0.0030), left medial orbitofrontal (R_(n=27)_ = -0.55; *p* = 0.0030), right medial orbitofrontal (R_(n=27)_ = -0.55, *p* = 0.0030), right rostral middle frontal (R_(n=27)_ = -0.55, *p* = 0.0030) and left frontal pole (R_(n=27)_ = -0.55, *p* = 0.0030).

## 4. Discussion

The findings of this study include decreased left amygdala volumes and frontal cortical thickness in schizophrenia and MAP compared to controls; increased volumes in bilateral caudate, putamen, and nucleus accumbens (NAcc) volumes in schizophrenia compared to controls; and increased right globus pallidus and bilateral NAcc volumes in schizophrenia compared to MAP. There were no significant differences across groups for cytokine levels and their associations with neuroimaging measures. Lastly, decreased left amygdala volumes and frontal cortical thickness were significantly associated with longer illness duration in schizophrenia and with longer duration of methamphetamine use in MAP.

The present data are consistent with previous findings that decreased left amygdala volumes and frontal cortical thinning are present in both schizophrenia ^40,41^ and MAP ^16,17^ compared to healthy controls, and are associated with duration of illness in schizophrenia ^42,43^. This study is the first to our knowledge showing decreased amygdala volume and cortical thickness in MAP. Taken together, these findings suggest important overlaps in the neurobiology of schizophrenia and MAP. Decreased cortical thickness and thus pyramidal neuron layers may reflect decreased neural density and functional dysconnectivity to subcortical and other cortical brain regions, which has been reported to underlie the symptoms of schizophrenia ^44–46^ and therefore may occur in MAP. However, further studies are needed to elucidate whether psychotic symptoms in MAP are indeed due to loss of neural connectivity from the frontal cortical neurons to subcortical and cortical structures.

Increased caudate volume was observed in schizophrenia but not in MAP, which is a consistent finding in schizophrenia and has been attributed to typical antipsychotic use ^6,47–50^. The finding of increased basal ganglia volumes has been replicated in several studies, including putamen volumes in chronic schizophrenia ^41,49,51–53^. The finding that globus pallidus and NAcc volume were increased in schizophrenia compared with MAP is a novel one, and may also highlight important differences in the neurobiology between schizophrenia and MAP. However, it is notable that increased globus pallidus volume was associated with increased duration of illness in schizophrenia, and that increased volume of these structures has been attributed to antipsychotic use. Given that use of antipsychotic agents is typically of lower duration in MAP, we cannot exclude the possibility that this finding points to differences in treatment between schizophrenia and MAP.

Decreased frontal cortical thickness was observed in the superior, middle and inferior frontal brain regions in schizophrenia compared to healthy controls, in addition to that of the left lateral OFC, widely reported finding in schizophrenia ^10,11,15,16,54–60^, this was also observed for the schizophrenia group who did not report methamphetamine use. Cortical thickness has been reported to decrease more rapidly throughout the course of the illness ^57,61^ compared to healthy controls and has also been observed in chronic schizophrenia ^14,62–64^. In MAP we observed significant decreases for the right medial OFC and lateral OFC cortical thickness compared to healthy controls, whilst the schizophrenia group showed significant decreases in cortical thickness of the lateral OFC and left medial OFC compared to MAP, suggesting a deleterious effect of schizophrenia on these regions in the left hemisphere. However, sensitivity analysis suggests that these findings were not observed in the schizophrenia group who reported no methamphetamine use, therefore this finding may be attributed to methamphetamine use instead of the structural differences we observed between the two psychotic disorders. This finding suggests decreased frontal cortical thickness in the right hemisphere in MAP compared to schizophrenia, occurring in the left hemisphere, which may aid in distinguishing MAP from schizophrenia. This is the first study to report decreased cortical thickness of the OFC regions in MAP, which has been reported in antipsychotic naïve schizophrenia individuals ^65^. We observed increased left frontal pole in the schizophrenia group compared to MAP as well as in the schizophrenia group who did not report methamphetamine use compared to those who reported methamphetamine use.

In MAP we observed significant decreases for the right lateral and medial OFC cortical thickness compared to healthy controls, whilst the schizophrenia group showed significant decreases in cortical thickness compared to that of MAP, particularly the lateral and left medial OFC. We speculate that the considerable duration of illness in the schizophrenia participants of this study may contribute to the differences between schizophrenia to MAP. In this respect, longer illness duration has been attributed to cortical thinning in patients with schizophrenia, including the PFC ^57,61^. Furthermore, increased duration of methamphetamine use was significantly associated with decreased frontal cortical thickness in MAP participants. Therefore, our results suggest a distinct deleterious effect of schizophrenia on the pathophysiology of reduced cortical thickness in the lateral and medial OFC regions in the left hemisphere.

We found no significant differences in cytokine concentrations across groups. Furthermore, no significant associations were observed between neuroimaging measures and cytokines levels. There is some inconsistency in the literature on immune markers on schizophrenia; our findings are consistent with some previous work ^67–71^. However, we cannot rule out the possibility that our negative findings are due to insufficient power. Moreover, the anti-inflammatory effects of antipsychotic medication in both schizophrenia and MAP groups may have ameliorated the inflammatory status in these study groups^70,72,73^.

A number of limitations deserve emphasis. First, the use of antipsychotics in both schizophrenia and MAP patients may have confounded the results reported in this study. Second, although our sample size is consistent with previous work on MAP, the small sample size warrants replication in a larger study. Third, self-reported methamphetamine abstinence was not supported by an objective confirmation. Fourth, sex differences were not examined due to the small sample of female participants. Fifth, methamphetamine use in some participants in the schizophrenia group may have confounded the comparisons with the MAP group, however our sensitivity analysis provides some reassurance that this is not the case.

### 4.1 Conclusions

This study shows distinctive structural variations in basal ganglia and amygdala volumes, in addition to frontal cortical thickness, in individuals with schizophrenia and MAP, suggesting that these structures play a pivotal role in the pathophysiology of these disorders. Differences in basal ganglia volumes between schizophrenia and MAP may be attributed by longer illness duration in schizophrenia. The novel discovery of increased globus pallidus and NAcc volumes in schizophrenia compared with MAP may show crucial distinctions in the neurobiology between these two conditions. To deepen our understanding of the neurobiological underpinnings of MAP, future investigations should employ larger sample sizes, incorporate longitudinal study designs, and integrate magnetic resonance spectroscopy which may show important neurometabolic signatures in these brain regions in MAP.

## Acknowledgements

We thank all study staff and staff at the Cape Universities Body Imaging Centre (CUBIC) of the University of Cape Town. We also wish to thank all who participated in this study.

## Funding

The study was supported by the Medical Research Council and National Research Foundation. L.B. also acknowledges her funding sources, administered by the University of Cape Town, National Research Foundation and Deutscher Akademischer Austauschdienst.

## Conflict of interest

**The authors have nothing to disclose.**

## Notes

### Competing Interest Statement

The authors have declared no competing interest.

### Summary of Updates

In MAP we observed significant decreases for the right medial OFC and lateral OFC cortical thickness compared to healthy controls, whilst the schizophrenia group showed significant decreases in cortical thickness of the lateral OFC and left medial OFC compared to MAP, suggesting a deleterious effect of schizophrenia on these regions in the left hemisphere. However, sensitivity analysis suggests that these findings were not observed in the schizophrenia group who reported no methamphetamine use, therefore this finding may be attributed to methamphetamine use instead of the structural differences we observed between the two psychotic disorders.

