## Supplementary File 1 for "Subcortical volumes, frontal cortical thickness, and pro-inflammatory cytokines in schizophrenia versus methamphetamine-induced psychosis"

Supplementary Table 1: Subcortical volume group differences across schizophrenia with and without methamphetamine use

| **Regions** | Effect size Cohen's d | CI: 95% | *p* | Effect size Hedges' g | t-value |
|---|---|---|---|---|---|
| Left hippocampus | 0.35 | [-0.35, 1.04] | 0.26 | 0.34 | 1.02 |
| Right hippocampus | 0.32 | [-0.38, 1.01] | 0.31 | 0.31 | 0.93 |
| Left amygdala | 0.22 | [-0.47, 0.91] | 0.50 | 0.21 | 0.64 |
| Right amygdala | 0.28 | [-0.41, 0.98] | 0.37 | 0.28 | 0.83 |
| Left caudate | 0.17 | [-0.53, 0.86] | 0.61 | 0.16 | 0.49 |
| Right caudate | 0.06 | [-0.63, 0.76] | 0.85 | 0.06 | 0.18 |
| Left putamen | 0.03 | [-0.66, 0.73] | 0.92 | 0.03 | 0.10 |
| Right putamen | 0.13 | [-0.57, 0.82] | 0.70 | 0.12 | 0.37 |
| Left globus pallidus | 0.18 | [-0.52, 0.87] | 0.59 | 0.17 | 0.52 |
| Right globus pallidus | 0.20 | [-0.50, 0.89] | 0.53 | 0.20 | 0.58 |
| Left nucleus accumbens | 0.06 | [-0.64, 0.75] | 0.86 | 0.06 | 0.17 |
| Right nucleus accumbens | 0.08 | [-0.61, 0.78] | 0.80 | 0.08 | 0.25 |

Effect size for all group differences. Parametric independent samples t-test performed for subcortical volumes in schizophrenia with methamphetamine use (SCZ_MA) and without methamphetamine use (SCZ), with *p*-values of $<0.05$ considered statistically significant. Effect sizes, *Cohens d at 95% confidence interval.

Supplementary Table 2: Frontal cortical thickness group differences across schizophrenia with and without methamphetamine use

| **Regions** | Effect size Cohen's d | CI: 95% | *p* | Effect size Hedges' g | t-value |
|---|---|---|---|---|---|
| **Left caudal middle frontal** | 0.56 | [-0.22, 1.35] | 0.13 | 0.55 | 1.46 |
| **Right caudal middle frontal** | 0.66 | [-0.22, 1.35] | 0.07 | 0.64 | 1.7 |
| **Right lateral orbitofrontal** | 0.71 | [-0.07, 1.49] | 0.045 | 0.69 | 1.84 |
| **Left rostral middle frontal** | 0.19 | [-0.59, 0.97] | 0.59 | 0.19 | 0.49 |
| **Left superior frontal** | 0.15 | [-0.63, 0.94] | 0.66 | 0.15 | 0.4 |
| **Right superior frontal** | 0.29 | [-0.49, 1.07] | 0.43 | 0.28 | 0.43 |
| **Left pars opercularis** | 0.54 | [-0.24, 1.32] | 0.15 | 0.53 | 1.4 |
| **Right pars opercularis** | 0.67 | [-0.11, 1.45] | 0.063 | 0.66 | 1.74 |
| **Left pars orbitalis** | 0.35 | [-0.43, 1.13] | 0.34 | 0.34 | 0.90 |
| **Right pars orbitalis** | 0.43 | [-0.35, 1.21] | 0.25 | 0.42 | 1.12 |
| **Left pars triangularis** | 0.07 | [-0.72, 0.85] | 0.85 | 0.06 | 0.17 |

Effect size for all group differences. Parametric independent samples t-test performed for frontal cortical thickness in schizophrenia with methamphetamine use (SCZ_MA) and without methamphetamine use (SCZ), with *p*-values of <0.05 considered statistically significant.. Effect sizes, *Cohens d at 95% confidence interval.

Supplementary Table 3: Frontal cortical thickness group differences across schizophrenia with and without methamphetamine use

| **Regions** | Biserial Correlation Coefficient *r* | CI: 95% | *p* | U |
|---|---|---|---|---|
| **Left lateral orbitofrontal** | 0.47 | [0.041, 0.34] | 0.039 | 64.5 |
| **Left medial orbitofrontal** | 0.44 | [0.054, 0.32] | 0.055 | 68.5 |
| **Right medial orbitofrontal** | 0.44 | [0.047, 0.33] | 0.050 | 67.5 |
| **Right rostral middle frontal** | 0.44 | [0.048, 0.33] | 0.054 | 68.5 |
| **Left frontal pole** | -0.47 | [0.064, -0.31] | 0.039 | 178.5 |
| **Right frontal pole** | 0.12 | [0.10, 0.28] | 0.6 | 106.5 |
| **Right pars triangularis** | 0.25 | [0.20, 0.24] | 0.27 | 91 |

Effect size for all group differences. Non-parametric Mann Whitney U performed for frontal cortical thickness in schizophrenia with methamphetamine use (SCZ_MA) and without methamphetamine use (SCZ), with *p*-values of <0.05 considered statistically significant. Effect sizes, *Biserial Correlation Coefficient ®
